# Seasonal decline in mitochondrial plasticity in the three-spined stickleback

**DOI:** 10.64898/2025.12.06.692738

**Authors:** Francisco Ruiz-Raya, Pat Monaghan, Caroline Millet, Neil B. Metcalfe

## Abstract

Mitochondrial plasticity enables ectothermic animals to maintain performance under changing thermal conditions, but whether this ability declines through adulthood is unknown. We explored changes in mitochondrial plasticity in the three-spined stickleback (*Gasterosteus aculeatus*), a temperate fish that in many populations has a single breeding season over which it produces multiple clutches and then dies. Wild-caught fish were exposed to one of three thermal regimes: remaining at 12 °C, switching to constant warm (21 °C) or diel cycling (12–21 °C) for three weeks at either the beginning or end of the season. We quantified both phosphorylating and non-ATP-producing (OXPHOS and LEAK) respiration in isolated muscle mitochondria, and oxidative control efficiency (OxCE). Early in the breeding season, young fish responded to elevated temperatures by adjusting both their capacity for ATP production and non-ATP-production respiration, but this mitochondrial plasticity declined over the season. OxCE was unaffected by time of season or temperature treatment. Changing body condition was unrelated to mitochondrial function. Our findings indicate that mitochondrial plasticity declines across adult life, potentially limiting thermal resilience in older individuals, the first time this has been reported in an ectotherm. This highlights the need to account for age-related physiological changes in capacity when predicting species’ responses to environmental change.

## 1. INTRODUCTION

Ectotherms show a limited capacity to use metabolic heat to maintain their body temperature, which makes them particularly vulnerable to thermal fluctuations [1,2]. However, they can adjust their physiology to buffer these fluctuations, a process known as thermal acclimation [1,3,4], so allowing them to limit the negative impacts of temperature variation and sustain performance under changing conditions [3,5,6]. Mitochondria are involved in this thermal acclimation due to their role in producing ATP [5–7]. By adjusting their respiratory capacity and the efficiency with which they produce energy in response to environmental temperature, mitochondria can influence thermal tolerance [8–11]. However, the potential for thermal plasticity in ectotherms has recently been shown to vary among taxa [12,13]. Limitations to this flexibility may reduce resilience and impair an animal’s performance and fitness [14,15]. Understanding the limits of mitochondrial plasticity is therefore crucial for predicting how species will cope with the increasing temperature fluctuations [16].

One potentially important, yet poorly understood, factor that might influence mitochondrial plasticity is age. As with other physiological traits, the capacity for thermal tolerance in ectotherms can vary across developmental stages but also decline with age [17]. Although the specific mechanisms underlying age-related changes in thermal tolerance are not fully understood, mitochondrial function generally decreases with advancing age [18–21], which raises the possibility that mitochondrial plasticity in response to temperature also diminishes over adult life. This may be particularly relevant for annual or short-lived species, where age-related reductions in mitochondrial performance could limit performance later in the single breeding season.

The three-spined stickleback (*Gasterosteus aculeatus*; hereafter “stickleback”) is a widespread teleost fish in the Northern Hemisphere [22], and a well-established model in eco-physiology and evolutionary biology [22,23]. Many freshwater stickleback populations are annual, with individuals producing multiple clutches but starting to senesce over a single breeding season, rarely surviving to a second year [24–26]. As the season advances, reproductive investment per clutch tends to increase while somatic condition declines, reflecting a shift towards a terminal investment [27,28]. Thus early-season adult fish are in their first reproductive stage with high somatic and physiological reserves (“young” within the season), while late-season individuals show signs of physiological ageing (“old” within the season) [27,29]. Sticklebacks show marked thermal acclimation capacity [14], linked to mitochondrial biology [30,31], but whether this mitochondrial plasticity declines over the course of the adults’ reproductive lifespan remains unknown.

Here, we experimentally tested whether mitochondrial thermal plasticity declines over the breeding lifetime in a freshwater population of three-spined sticklebacks. Early- and late-season adult females were exposed to either constant cool (12°C), constant warm (21°C), or diel cycling (12–21°C) temperatures – the latter treatment better mimicking the cycles in water temperature that populations experience within a single day [32,33]. Mitochondrial respiratory capacity was quantified as the oxygen consumption associated with either ATP production (i.e., oxidative phosphorylation; OXPHOS) or offsetting the leak of protons across the inner mitochondrial membrane (i.e., LEAK, which does not generate ATP). This allowed us to calculate mitochondrial efficiency, which we define as the proportion of oxygen consumption contributing to oxidative phosphorylation (Oxidative Control Efficiency, OxCE). We predicted that early-season (“young adult”) females would show higher mitochondrial plasticity (i.e., higher variation in OXPHOS and LEAK rates among temperature treatments) than end-of-season (“old adult”) individuals, reflecting declines in acclimation potential associated with senescence. Diel cycling temperatures are predicted to elicit intermediate mitochondrial responses compared to constant thermal regimes. Because thermal acclimation can affect energy balance [34], we also measured body condition as a proxy for whole-organism state [35], and used structural equation modelling (SEM) to test whether any temperature effects on body condition were mediated by mitochondrial function.

## 2. METHODS

### Study species and fish husbandry

Pre-reproductive adult three-spined sticklebacks (*Gasterosteus aculeatus*) were collected in early March 2024 from the River Kelvin, Glasgow, UK (55°52’29.1”N, 4°16’48.5”W) using dip nets and minnow traps. Stream populations of sticklebacks at mid-latitudes (50–60° N), such as our study population, generally reproduce for 3.5–4 months during spring and summer (April–August) [36]. Captured fish (N = 200) were transported to the laboratory where they were housed in groups in 80-L tanks with shelter and recirculating UV-sterilized water maintained at 12 °C (the approximate water temperature of Scottish rivers at the start of the breeding season; see below) and a 12:12 light:dark photoperiod. To reduce social stress, males exhibiting breeding coloration were removed, leaving only females and nonreproductive males. Females could still lay clutches of eggs, which were not reared and degraded over time; however, any energetic costs of egg production were likely to be minor due to ad libitum feeding. Fish were fed twice daily with commercial pellet food. Full details of fish collection, transport, and housing are provided in the Supplementary Material.

### Thermal manipulation

The thermal acclimation trials were designed to allow us to examine whether protracted exposure (3 weeks) or diel cycling exposure to high temperatures changed mitochondrial function. We conducted the trials on young females in April just before the commencement of the breeding season, and on older females after the breeding season was over (September) but before any significant mortality had occurred (the majority of the fish, 84% in the holding tanks, survived to the end of season sampling date). In both age groups, 63 female fish from the holding tanks were randomly assigned to one of three temperature regimes for 21 days: no change (i.e. temperature remains the same as the holding tank 12°C; *n* = 21), warmer temperature (increased to 21°C; *n* = 21), or diel cycling 12–21°C; *n* = 21). Temperatures were chosen according to previous research [37–39], and based on spring–summer river temperature records from Scotland in 2023, which averaged 13.1 ± 1.2 °C (mean ± SD) and typically ranged between 8.2 ± 1.9 and 20.2 ± 2.2 °C [40]. Adult three-spined sticklebacks show a critical thermal maximum (CTmax) of approximately 33 °C [41], indicating that our upper treatment of 21°C represents a warm but ecologically relevant condition.

Since mitochondrial measurements at the end of the 21-day temperature trial could only be performed on seven fish per day, the experimental tanks were populated with fish from the holding tanks at a rate of 1 experimental tank per day (rotating across the three treatments to prevent sequence effects). Each temperature regime was replicated across three experimental tanks (27 × 48 × 36 cm; 47 L), with seven fish per tank, for a total of nine tanks. All tanks were housed in the same room, with a constant photoperiod (12L:12D). The water temperature in the tanks was controlled using programmable heaters. For the diel cycling treatment, heaters were placed on a timer to generate the diel cycle (12–21 °C; see Figure S1 for temperature traces). Fish were fed *ad libitum* twice daily on weekdays and once daily on weekends using commercial pellet food.

### Tissue sampling and mitochondria isolation

After the 21-day temperature trial, all fish within a tank were euthanised (using a benzocaine overdose of 10 mg/L) for measurements of mitochondrial function. Each individual was blotted dry, and its mass recorded (to 0.001g). The total length (including the caudal fin) of each fish was measured (to 0.1mm) using vernier callipers. Isolates of muscle mitochondria were prepared using a modified version of previous protocols [42]. Briefly, immediately after death, muscle samples from the myotome were collected and placed on ice in isolation buffer; these were minced and gently homogenized using a Teflon-on-glass homogenizer with six strokes at 100 rpm. The resulting homogenate was centrifuged at 1000 g (10 min). The supernatant was then centrifuged at 8700 g (10 min), and resulting pellets were resuspended in in 1000 µl of storage buffer. A portion of the mitochondrial preparation was kept on ice for immediate measurement of mitochondrial function, and the remainder was stored at −80 °C for subsequent enzymatic analysis (see Supplementary Material for a detailed description of mitochondrial isolation methods).

### Mitochondrial function measurements

Mitochondrial function was assessed at 21 °C using high-resolution respirometry on Oxygraph-2k systems (Oroboros Instruments, Innsbruck, Austria). Isolated mitochondrial samples were introduced into the respirometry chamber, and oxygen consumption was determined by monitoring the rate of oxygen decline in the sealed chamber after the addition of substrates (see Supplementary Material). The rate of respiration in the presence of all substrates represents the maximum rate of OXPHOS, which can be used as a proxy for the capacity for mitochondrial ATP production. We determined LEAK respiration, being the oxygen consumption that is used to offset proton leak across the inner mitochondrial membrane rather than to generate ATP. We also measured the increase in respiration after addition of exogenous cytochrome C (CytC) as a quality-control test for outer membrane permeability. Any sample showing compromised quality was excluded (see Supplementary Material). For the fish kept continuously at 12 °C, mitochondrial respiration was measured at both 12 °C and 21 °C to evaluate whether assay temperature influenced potential age-related effects on mitochondrial function.

Respiration rates were normalized to citrate synthase (CS) activity, a widely used biomarker for the mitochondrial content in the assayed sample [43,44]. This method allowed us to control for differences in mitochondrial volume density among preparations. All CS assays were performed under identical, temperature-controlled conditions, ensuring that normalization preserves relative differences among samples (see Supplementary Material for a detailed description of CS activity measurements).

Mitochondrial Oxidative Control Efficiency (OxCE; sometimes also referred as OXPHOS Coupling Efficiency [11]) was calculated as: OxCE = 1 - (LEAK / OXPHOS). This mitochondrial efficiency index ranges from hypothetical values of 0 (indicating no ATP production despite substrate availability) to 1 (indicating absence of leak respiration and all oxygen consumption dedicated to ATP synthesis). Overall, respiration rates indicate potential metabolic capacity levels, while OxCE indicates how efficient mitochondria are at using oxygen when producing ATP [11].

### Statistical analysis

All statistical analyses were conducted in R version 4.3.3 [45]. We analysed changes in mitochondrial capacity (i.e., OXPHOS and LEAK respiration) and efficiency (OxCE) using separate generalized linear mixed models (GLMMs) implemented in the *glmmTMB* package [46]. All models included temperature treatment (12°C, 21°C, or 12-21°C), age (young adult vs. old adult), and their interaction, as fixed factors. CytC response was also included as a covariate to control for potential effects of mitochondria outer membrane permeability. Experimental tank ID and respirometry chamber ID were included as random factors in all models. Body condition index (BCI) was calculated as the residuals from an OLS regression of log(mass) against log(body total length), with age included as an additional predictor. Variation in BCI was analysed using GLMM with the same fixed and random structure described above. Normality, homoscedasticity and singularity assumptions were evaluated using the *performance* package [47], and models with singular fits were refitted with specified Gamma priors [48]. *P*-values were obtained from Wald statistics by using the *car* package [49]. Post hoc pairwise comparisons and standardized effect sizes (Cohen’s d and 95% CI) were obtained using the *emmeans* package [50]. All values in plots are presented as estimated marginal means ± 95% CI.

To test whether the effects of temperature treatment on body condition were mediated by changes in mitochondrial function, we conducted a structural equation model (SEM) using the *piecewiseSEM* package [51]. We explored the direct link between temperature treatment and body condition, as well as the indirect path through changes in mitochondrial function. SEM was restricted to young adults, as mitochondrial function was not affected by temperature treatment in old individuals (see Results section). OXPHOS and LEAK were modelled in separate LMM (*glmmTMB* package) as a function of temperature treatment and CytC. Body condition was modelled using LMMs (*glmmTMB* package), incorporating temperature treatment and either OXPHOS or LEAK as predictors in separate models. Tank ID and respirometry chamber ID were included as random effects in all models to account for potential experimental grouping effects. Multicollinearity among predictors was assessed via variance inflation factors (VIFs; all cases VIF< 2) using the *performance* package [52]. We present the SEM including OXPHOS as the mediator in the main text, while results with LEAK as the mediator are provided in the Supplementary Material.

## 3. RESULTS

Temperature treatment significantly influenced OXPHOS respiration, but this effect was restricted to young adults (temperature × age interaction: χ^2^ = 23.92; df = 2; *p* < 0.001; Figure 1a). Young fish showed marked variation across temperature regimes, with lower OXPHOS rates in the constant warm (21 °C) group compared with young adults kept at a constant cool temperature (12 °C; estimate = −2.36; SE = 0.66; *p* = 0.009). OXPHOS rates under diel cycling temperatures were intermediate and did not differ significantly from the constant cool and constant warm groups (all *p* > 0.15). In contrast, OXPHOS respiration did not vary among temperature treatments in the females tested later in the season (all *p* > 0.30), indicating reduced mitochondrial thermal plasticity in the older females. Notably, old adults kept at 21 °C had higher OXPHOS rates than fish in the same temperature treatment tested earlier in the season (estimate = 1.53; SE = 0.44; *p* = 0.010).

**Figure 1.**
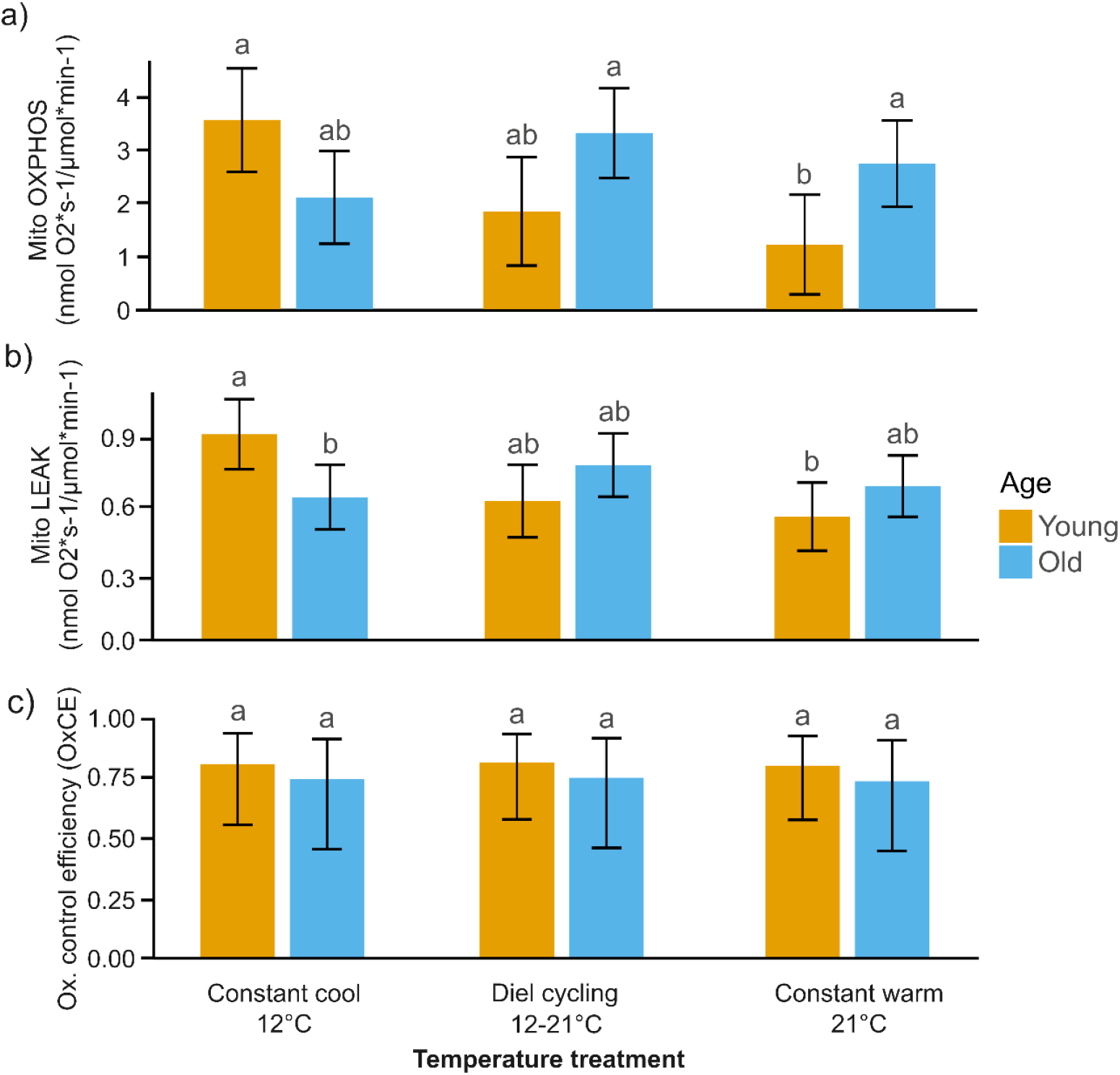
Effects of temperature treatment on mitochondrial capacity and efficiency in young adult vs. old adult three-spined stickleback females. (a) ATP-producing respiration (OXPHOS), (b) non-ATP-producing respiration (LEAK), and (c) oxidative control efficiency (OxCE) after exposure to constant cool (12 °C), diel cycling (12–21 °C), and constant warm (21 °C) treatments. Bars represent means ± 95% CI for young (orange; *n* = 9, 9, and 10 for cool, diel cycling, and warm, respectively) and old (blue; *n* = 14, 17, and 18 for cool, diel cycling, and warm, respectively) females. Different letters indicate statistically significant differences among the six treatment groups (Tukey post hoc tests, *p* < 0.05).

LEAK respiration differed between young and old fish depending on temperature treatment, as indicated by a significant age × temperature interaction (χ^2^ = 23.82; df = 2; *p* < 0.0001; Figure 1b), even though there was no overall effect of age (χ^2^ = 0.193; df = 1; *p* = 0.661). In young females, LEAK was significantly lower in the constant warm (21 °C) treatment than the constant cool (12 °C) regime (Tukey = 0.37; SE = 0.11; *p* = 0.012), consistent with a temperature-induced reduction in proton leak. LEAK rates in the diel cycling treatment were intermediate between the constant cool and constant warm treatments (all *p* > 0.09). In contrast, older individuals showed no significant variation in LEAK respiration across temperature treatments (all *p* > 0.66).

In the fish kept at a constant cool (12 °C) temperature and whose mitochondrial function was measured at both assay temperatures (12 °C and 21°C), there was no age × assay-temperature interaction for OXPHOS (χ^2^ = 0.60, *p* = 0.437) or LEAK respiration (χ^2^ = 0.20, *p* = 0.654), meaning that age differences were not influenced by assay temperatures (Figure S2). Finally, no significant effects of the temperature treatment (χ^2^ = 0.01; df = 2; *p* = 0.994; Figure 1c), age (χ^2^ = 0.37; df = 1; *p* = 0.542), or CytC content (χ^2^ = 0.35; df = 1; *p* = 0.552) on oxidative control efficiency (OxCE).

We found a significant effect of temperature treatment on body condition (χ^2^ = 6.28; df = 2; *p* = 0.043), with fish in the constant warm regime showing lower BCI. The age × temperature interaction was not statistically significant (χ^2^ = 5.25, df = 2, *p* = 0.072), but effect sizes indicated age-specific trends. Younger fish showed markedly lower BCI under constant warm conditions relative to the constant-cool treatment (Cohen’s d = 1.56, 95% CI [0.55, 2.57]). In older fish, temperature effects were consistently small (Cohen’s d < 0.4). Age alone had no significant effect on BCI (χ^2^ = 0.002; df = 1; *p* = 0.969).

Our structural equation modelling (SEM) confirmed that temperature treatments significantly affected both OXPHOS capacity and body condition (BCI) in young adults (Figure 2b; Table S3). However, no significant link was found between OXPHOS and BCI (Figure 2b; Table S2), indicating that variation in mitochondrial function did not drive changes in BCI. Specifically, young fish kept at the constant cool temperature (12 °C) showed a significant increase in OXPHOS capacity (estimate = 0.876; *p* = 0.012; Table S3), but this was not linked to changes in BCI. Conversely, fish in the constant warm (21 °C) treatment exhibited a significant reduction in BCI (estimate = –0.831; *p* = 0.005; Table S3), yet this decline was not mediated by OXPHOS capacity.

**Figure 2.**
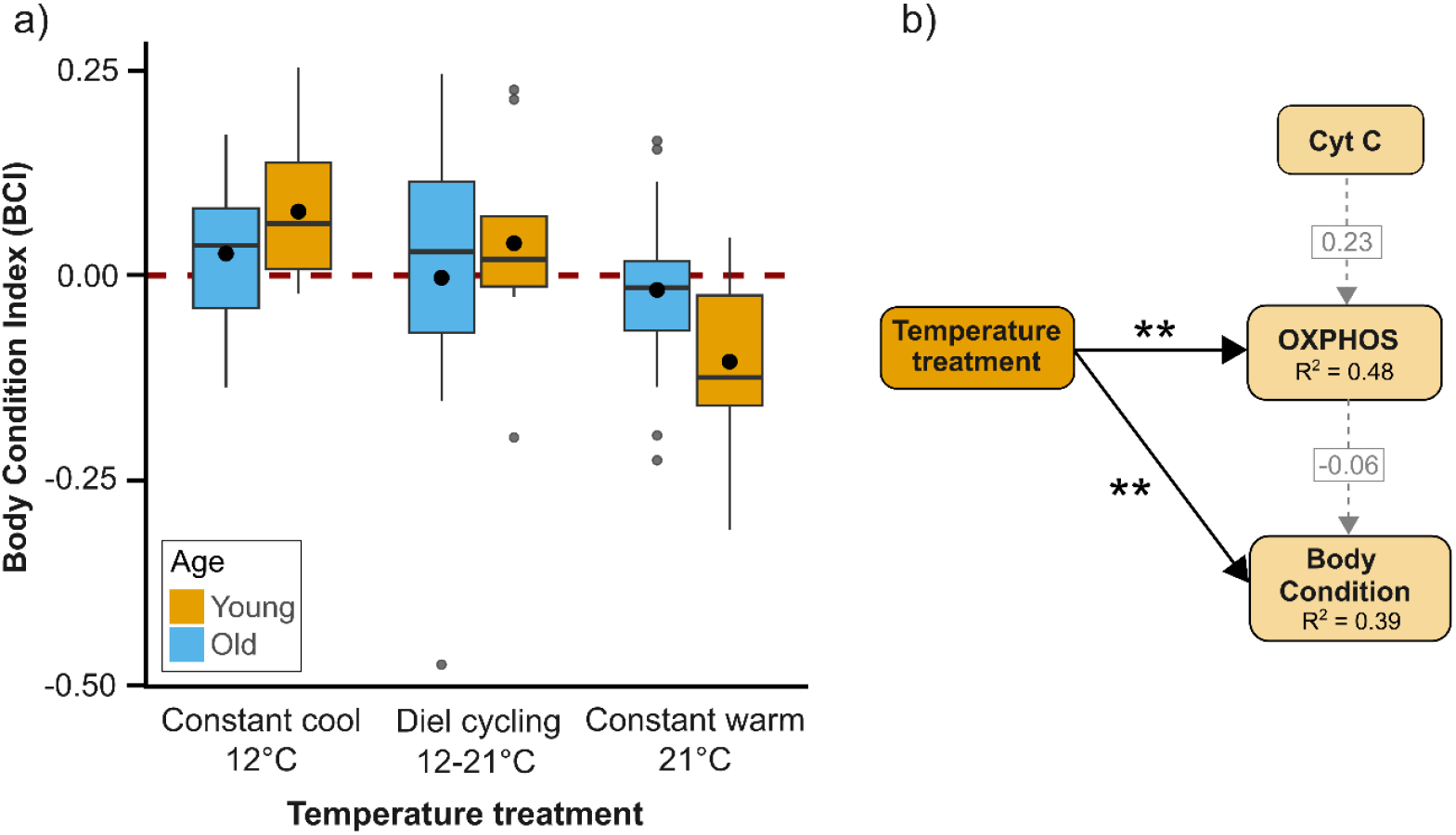
a) Effects of temperature treatment and age on body condition in fish. Body condition index (BCI) is shown for young (orange) and old (blue) adults across three temperature treatments: constant cool (12°C), diel cycling (12–21°C), and constant warm (21°C). Bars represent means ± 95% CI. b) Structural equation models showing direct and indirect paths between temperature treatment and BCI. Indirect paths included mitochondrial capacity for ATP production (OXPHOS). Grey dotted arrows represent non-significant paths, while black solid lines represent significant paths (\*\**p* < 0.01). *R*^2^ represents the proportion of variance explained in response variables by a set of predictors. Standardized estimates are reported for those paths including continuous variables.

Similarly, we found no evidence that LEAK respiration mediated the effects of temperature treatment on BCI: while thermal treatment significantly affected both LEAK respiration and BCI, LEAK showed no significant association with BCI (Table S4; Figure S3).

## 4. DISCUSSION

Thermal acclimation is well documented across ectotherm taxa [12,53–55], an attribute that could deteriorate with age [17]. One potential mechanism underlying such age-related deterioration might be reduced flexibility in mitochondrial function with age, since mitochondria play a crucial role in thermal tolerance as the primary site of ATP production [56,57]. Our results show that mitochondrial plasticity declines with adult age in the three-spined stickleback. Specifically, young adult females at the start of their breeding season showed the capacity to make thermal-induced adjustments in both their phosphorylating (OXPHOS) and non-ATP-producing (LEAK) respiration when exposed to warm conditions, indicating flexibility in mitochondrial respiratory capacity. Responses in the diel cycling treatment were overall intermediate between the constant cool and warm regimes, indicating partial acclimation to diel thermal variability. Old adults, in contrast, showed limited mitochondrial responsiveness to temperature, suggesting that this plasticity diminishes as individuals age. These findings provide the first evidence, to our knowledge, that mitochondrial thermal plasticity declines with age in any ectotherm.

Such a decline in mitochondrial plasticity might reflect rapid age-related deterioration in sticklebacks, which is typically accompanied by accumulated reproductive costs and accelerated somatic decline [27]. At the mitochondrial level, this aging may involve accumulated mtDNA damage, loss of protein quality control, and an altered balance between mitochondrial fusion and fission, resulting in reduced energy production and elevated reactive oxygen species (ROS) [57]. While mild mitochondrial stress can activate protective responses that sustain function, these responses may weaken with age [18–21], potentially narrowing the range over which mitochondria can adjust to temperature changes. However, it is important to note that, although our cross-sectional design (necessitated by the terminal sampling required to measure mitochondrial function) cannot fully distinguish between within-individual ageing and selective disappearance of less plastic individuals, the low mortality between sampling points (∼16%) suggests that selective disappearance could have played only a minor role. Finally, data from the young and old fish that did not experience any temperature elevation, whose mitochondrial function was measured at both 12 °C and 21 °C, indicate that age-related differences are consistent across assay temperatures.

Although mitochondrial capacity for ATP production was less plastic in the older sticklebacks, mitochondrial oxidative control efficiency (i.e., the proportion of oxygen contributed to oxidative phosphorylation rather than lost to offset the proton leak) did not differ between the young and old females and was unaffected by thermal regime. Since this determines how much of the assimilated resources can be allocated to life-history traits and whole-organism performance [11], such preserved function might suggest that mitochondria become more stable in performance over the season, helping older fish to buffer the effect on performance in variable thermal environments. In our study, all fish were maintained under constant laboratory conditions, including a fixed photoperiod, which means that observed differences could be attributed to physiological aging rather than environmental changes. However, this protocol results in the exclusion of natural photoperiodic cues, which can influence reproductive timing and metabolic adjustments in the wild [58]. Similarly, variation in egg production, which it was not possible to measure in this study since eggs laid in the tanks could not be attributed to individual females, may also play a role.

Importantly, we also found that a relatively brief (3-week) exposure to warmer temperatures (21 °C) led to a decrease in individual body condition, particularly in young adults. This pattern matched our mitochondrial results: younger adults, which showed greater mitochondrial plasticity, may be more able to cope with higher energetic costs that divert resources from somatic maintenance. In contrast, older fish, with reduced mitochondrial plasticity but stable efficiency, may avoid these costs, helping to preserve body condition. However, our SEM results provided no evidence that mitochondrial function directly mediated these effects. Differences between young and old adults may also reflect age-related shifts in energy allocation, with younger fish investing more in growth and reproductive maturation, while older individuals conserve energy through reduced activity or altered resource use. Although reduced mitochondrial plasticity in older adults did not translate into lower mitochondrial efficiency or poorer condition under the conditions tested here, it remains possible that limited plasticity could constrain acclimation under more extreme or rapidly fluctuating temperatures.

In conclusion, we show that young sticklebacks at the start of their breeding season have higher mitochondrial plasticity than do older conspecifics at the end of the breeding season. These findings have potential consequences for population resilience under increasing climatic variability [1]. It is currently unknown whether similar patterns occur in other species or ecological contexts. What is now needed are longitudinal studies that examine multiple tissues (e.g., liver, heart) and additional hallmarks of aging (e.g., ROS production, mitochondrial DNA integrity). Such work will be critical for linking changes in mitochondrial function to ecological outcomes, as well as for assessing whether plasticity can be maintained or restored as climate change increases the pace and unpredictability of temperature variation in natural systems.

## Ethics

All animal husbandry and subsequent experiments were undertaken under UK Home Office project license PP6899400.

## Data availability statement

Datasets generated during this study are available from the corresponding author on reasonable request.

## Authors’ contributions

F.R.R.: conceptualization, funding acquisition, project administration, methodology, investigation, software, data curation, formal analysis, visualization, writing—original draft; P.M.: conceptualization, funding acquisition, project administration, resources, supervision, methodology, writing—review and editing; C.M.: investigation, writing—review and editing; N.B.M.: conceptualization, funding acquisition, project administration, resources, supervision, methodology, writing—review and editing. All authors gave final approval for publication and agreed to be held accountable for the work performed therein.

## Conflict of interest declaration

We declare we have no competing interests.

## Funding

This research was funded by a Marie Skłodowska-Curie Fellowship (HORIZON-MSCA-2022-PF-01; UKRI Guarantee fund EP/Y009908/1) awarded to FRR, and an ERC Advanced Grant (834653) awarded to NBM. PM was supported by an ERC Advanced Grant from the European Union’s Horizon 2020 research and innovation programme (grant agreement No. 101020037 to PM).

## Acknowledgements

We thank the members of Biological Services at the University of Glasgow, particularly Ross Phillips, Catherine Law, Catriona Mills, and Joseph Pennock, for their assistance with experimental setup and fish husbandry. We are also grateful to Darryl McLennan for help with fish collection and husbandry, and to Neal J. Dawson for his advice on laboratory analyses.

## Notes

### Competing Interest Statement

The authors have declared no competing interest.

## REFERENCES

1. Seebacher F, White CR, Franklin CE. 2015 Physiological plasticity increases resilience of ectothermic animals to climate change. Nat Clim Chang 5, 61–66. (doi:10.1038/NCLIMATE2457;SUBJMETA=158,2455,631;KWRD=ECOPHYSIOLOGY)

2. Bickford DP, Sheridan JA, Howard SD. 2011 Climate change responses: Forgetting frogs, ferns and flies? Trends Ecol Evol 26, 553–554. (doi:10.1016/j.tree.2011.06.016)

3. Angilletta MJ, Bennett AF, Guderley H, Navas CA, Seebacher F, Wilson RS. 2006 Coadaptation: A unifying principle in evolutionary thermal biology. Physiological and Biochemical Zoology 79, 282–294. (doi:10.1086/499990/ASSET/IMAGES/LARGE/FG6.JPEG)

4. Hoffmann AA, Sgró CM. 2011 Climate change and evolutionary adaptation. Nature 470, 479–485. (doi:10.1038/NATURE09670;SUBJMETA=181,457,631,649,704,841,844;KWRD=CLIMATE-CHANGE+ADAPTATION,GENETIC+VARIATION)

5. Iftikar FI, Hickey AJR. 2013 Do mitochondria limit hot fish hearts? Understanding the role of mitochondrial function with heat stress in notolabrus celidotus. PLoS One 8, e64120. (doi:10.1371/JOURNAL.PONE.0064120)

6. Pörtner H. 2001 Climate change and temperature-dependent biogeography: Oxygen limitation of thermal tolerance in animals. Naturwissenschaften 88, 137–146. (doi:10.1007/S001140100216/METRICS)

7. Camus MF, Wolff JN, Sgrò CM, Dowling DK. 2017 Experimental support that natural selection has shaped the latitudinal distribution of mitochondrial haplotypes in Australian Drosophila melanogaster. Mol Biol Evol 34, 2600–2612. (doi:10.1093/MOLBEV/MSX184)

8. Fangue NA, Richards JG, Schulte PM. 2009 Do mitochondrial properties explain intraspecific variation in thermal tolerance? Journal of Experimental Biology 212, 514–522. (doi:10.1242/JEB.024034)

9. Pichaud N, Ekström A, Breton S, Sundström F, Rowinski P, Blier PU, Sandblom E. 2019 Cardiac mitochondrial plasticity and thermal sensitivity in a fish inhabiting an artificially heated ecosystem. Sci Rep 9, 1–11. (doi:10.1038/S41598-019-54165-3;SUBJMETA=1465,1737,319,333,443,601,631;KWRD=ANIMAL+PHYSIOLOGY,ENERGY+METABOLISM)

10. Chung DJ, Morrison PR, Bryant HJ, Jung E, Brauner CJ, Schulte PM. 2017 Intraspecific variation and plasticity in mitochondrial oxygen binding affinity as a response to environmental temperature. Sci Rep 7, 1–10. (doi:10.1038/S41598-017-16598-6;SUBJMETA=158,1737,2455,601,631;KWRD=ANIMAL+PHYSIOLOGY,ECOPHYSIOLOGY)

11. Koch RE et al. 2021 Integrating mitochondrial aerobic metabolism into ecology and evolution. Trends Ecol Evol 36, 321–332. (doi:10.1016/J.TREE.2020.12.006/ASSET/0EBF99B9-96F2-40DC-B316-EF6FE592DB20/MAIN.ASSETS/MMC2.MP4)

12. Burton T, Einum S. 2025 High Capacity for Physiological Plasticity Occurs at a Slow Rate in Ectotherms. Ecol Lett 28, e70046. (doi:10.1111/ELE.70046)

13. Einum S, Burton T. 2022 Divergence in rates of phenotypic plasticity among ectotherms. Ecol Lett 26, 147–156. (doi:10.1111/ELE.14147;PAGEGROUP:STRING:PUBLICATION)

14. De Bonville J et al. 2025 Dynamics of thermal tolerance plasticity across fish species and life stages. J Therm Biol 127, 104024. (doi:10.1016/J.JTHERBIO.2024.104024)

15. Sandblom E et al. 2016 Physiological constraints to climate warming in fish follow principles of plastic floors and concrete ceilings. Nat Commun 7, 1–8. (doi:10.1038/NCOMMS11447;SUBJMETA=158,1737,2165,2722,601,631;KWRD=ANIMAL+PHYSIOLOGY,CLIMATE-CHANGE+ECOLOGY,ICHTHYOLOGY)

16. Diffenbaugh NS, Field CB. 2013 Changes in ecologically critical terrestrial climate conditions. Science (1979) 341, 486–492. (doi:10.1126/SCIENCE.1237123/ASSET/22D14C23-8FC0-47F9-A1DE-6FAED6838E8C/ASSETS/GRAPHIC/341_486_F4.JPEG)

17. Bowler K, Terblanche JS. 2008 Insect thermal tolerance: What is the role of ontogeny, ageing and senescence? Biological Reviews 83, 339–355. (doi:10.1111/J.1469-185X.2008.00046.X;SUBPAGE:STRING:FULL)

18. Short KR, Bigelow ML, Kahl J, Singh R, Coenen-Schimke J, Raghavakaimal S, Nair KS. 2005 Decline in skeletal muscle mitochondrial function with aging in humans. Proc Natl Acad Sci U S A 102, 5618–5623. (doi:10.1073/PNAS.0501559102;WGROUP:STRING:PUBLICATION)

19. Amara CE, Shankland EG, Jubrias SA, Marcinek DJ, Kushmerick MJ, Conley KE. 2007 Mild mitochondrial uncoupling impacts cellular aging in human muscles in vivo. Proc Natl Acad Sci U S A 104, 1057–1062. (doi:10.1073/PNAS.0610131104/SUPPL_FILE/10131FIG5.JPG)

20. Salmón P, Dawson NJ, Millet C, Selman C, Monaghan P. 2023 Mitochondrial function declines with age within individuals but is not linked to the pattern of growth or mortality risk in zebra finch. Aging Cell 22, e13822. (doi:10.1111/ACEL.13822;JOURNAL:JOURNAL:14749726;PAGE:STRING:ARTICLE/CHAPTER)

21. Grevendonk L et al. 2021 Impact of aging and exercise on skeletal muscle mitochondrial capacity, energy metabolism, and physical function. Nature Communications 2021 12:1 12, 1–17. (doi:10.1038/s41467-021-24956-2)

22. McKinnon JS, Rundle HD. 2002 Speciation in nature: The threespine stickleback model systems. Trends Ecol Evol 17, 480–488. (doi:10.1016/S0169-5347(02)02579-X/ASSET/15406448-9B62-4004-BAFA-6A330753F0C8/MAIN.ASSETS/GR1.JPG)

23. Pottinger TG, Carrick TR, Yeomans WE. 2002 The three-spined stickleback as an environmental sentinel: Effects of stressors on whole-body physiological indices. J Fish Biol 61, 207–229. (doi:10.1111/J.1095-8649.2002.TB01747.X;WGROUP:STRING:PUBLICATION)

24. Lee WS, Monaghan P, Metcalfe NB. 2012 The pattern of early growth trajectories affects adult breeding performance. Ecology 93, 902–912. (doi:10.1890/11-0890.1;SUBPAGE:STRING:FULL)

25. Baker JA, Wund MA, Heins DC, King RW, Reyes ML, Foster SA. 2015 Life-history plasticity in female threespine stickleback. Heredity (Edinb) 115, 322–334. (doi:10.1038/HDY.2015.65;SUBJMETA=181,631;KWRD=EVOLUTION)

26. Baker JA, Heins DC, Foster SA, King RW. 2008 An overview of life-history variation in female three-spine stickleback. Behaviour 145, 579–602.

27. Poizat G, Rosecchi E, Crivelli AJ. 1999 Empirical evidence of a trade–off between reproductive effort and expectation of future reproduction in female three-spined sticklebacks. Proc R Soc Lond B Biol Sci 266, 1543–1548. (doi:10.1098/RSPB.1999.0813)

28. Magierecka A, Aristeidou A, Papaevripidou M, Gibson JK, Sloman KA, Metcalfe NB. 2022 Timing of reproduction modifies transgenerational effects of chronic exposure to stressors in an annual vertebrate. Proceedings of the Royal Society B: Biological Sciences 289. (doi:10.1098/RSPB.2022.1462;PAGE:STRING:ARTICLE/CHAPTER)

29. Álvarez-Quintero N, Velando A, Noguera JC, Kim SY. 2020 Environment-induced changes in reproductive strategies and their transgenerational effects in the three-spined stickleback. Ecol Evol 11, 771. (doi:10.1002/ECE3.7052)

30. Cominassi L, Ressel KN, Brooking AA, Marbacher P, Ransdell-Green EC, O’Brien KM. 2022 Metabolic rate increases with acclimation temperature and is associated with mitochondrial function in some tissues of threespine stickleback. Journal of Experimental Biology 225. (doi:10.1242/JEB.244659/278444/AM/METABOLIC-RATE-INCREASES-WITH-ACCLIMATION)

31. Keenan K, Hoffman M, Dullen K, O’Brien KM. 2017 Molecular drivers of mitochondrial membrane proliferation in response to cold acclimation in threespine stickleback. Comp Biochem Physiol A Mol Integr Physiol 203, 109–114. (doi:10.1016/J.CBPA.2016.09.001)

32. Corey E, Linnansaari T, Cunjak RA, Currie S. 2017 Physiological effects of environmentally relevant, multi-day thermal stress on wild juvenile Atlantic salmon (Salmo salar). Conserv Physiol 5, cox014. (doi:10.1093/CONPHYS/COX014)

33. Broadmeadow SB, Jones JG, Langford TEL, Shaw PJ, Nisbet TR. 2011 The influence of riparian shade on lowland stream water temperatures in southern England and their viability for brown trout. River Res Appl 27, 226–237. (doi:10.1002/RRA.1354)

34. Sandblom E, Gräns A, Axelsson M, Seth H. 2014 Temperature acclimation rate of aerobic scope and feeding metabolism in fishes: Implications in a thermally extreme future. Proceedings of the Royal Society B: Biological Sciences 281. (doi:10.1098/RSPB.2014.1490;WGROUP:STRING:PUBLICATION)

35. Peig J, Green AJ. 2009 New perspectives for estimating body condition from mass/length data: the scaled mass index as an alternative method. Oikos 118, 1883–1891. (doi:10.1111/j.1600-0706.2009.17643.x)

36. Ishikawa A, Kitano J. 2020 Diversity in reproductive seasonality in the three-spined stickleback, Gasterosteus aculeatus. Journal of Experimental Biology 223. (doi:10.1242/JEB.208975,)

37. Jackson FL, Fryer RJ, Hannah DM, Millar CP, Malcolm IA. 2018 A spatio-temporal statistical model of maximum daily river temperatures to inform the management of Scotland’s Atlantic salmon rivers under climate change. Science of The Total Environment 612, 1543–1558. (doi:10.1016/J.SCITOTENV.2017.09.010)

38. Morris MRJ, Wuitchik SJS, Rosebush J, Rogers SM. 2021 Mitochondrial volume density and evidence for its role in adaptive divergence in response to thermal tolerance in threespine stickleback. J Comp Physiol B 191, 657–668. (doi:10.1007/S00360-021-01366-W/METRICS)

39. Magierecka A, Lind ÅJ, Aristeidou A, Sloman KA, Metcalfe NB. 2021 Chronic exposure to stressors has a persistent effect on feeding behaviour but not cortisol levels in sticklebacks. Anim Behav 181, 71–81. (doi:10.1016/J.ANBEHAV.2021.08.028)

40. Scottish Government. 2025 Scotland River Temperature Monitoring Network (SRTMN): Outputs and tools. See https://www.gov.scot/publications/scotland-river-temperature-monitoring-network-srtmn/pages/outputs-and-tools/ (accessed on 11 September 2025).

41. Cowan ZL, Andreassen AH, De Bonville J, Green L, Binning SA, Silva-Garay L, Jutfelt F, Sundin J. 2023 A novel method for measuring acute thermal tolerance in fish embryos. Conserv Physiol 11, coad061. (doi:10.1093/CONPHYS/COAD061)

42. Dawson NJ, Scott GR. 2022 Adaptive increases in respiratory capacity and O2 affinity of subsarcolemmal mitochondria from skeletal muscle of high-altitude deer mice. The FASEB Journal 36, e22391. (doi:10.1096/FJ.202200219R)

43. McLaughlin KL, Hagen JT, Coalson HS, Nelson MAM, Kew KA, Wooten AR, Fisher-Wellman KH. 2020 Novel approach to quantify mitochondrial content and intrinsic bioenergetic efficiency across organs. Sci Rep 10, 1–15. (doi:10.1038/S41598-020-74718-1;SUBJMETA=1465,319,333,443,45,475,631;KWRD=ENERGY+METABOLISM,PROTEOMICS)

44. Larsen S et al. 2012 Biomarkers of mitochondrial content in skeletal muscle of healthy young human subjects. J Physiol 590, 3349–3360. (doi:10.1113/JPHYSIOL.2012.230185)

45. R Core Team. 2022 R: A language and environment for statistical computing. R Foundation for Statistical Computing, Vienna, Austria. URL https://www.R-project.org/.

46. Brooks M, Kristensen K, van Benthe K, Magnusson A, Berg C, Nielsen A, Skaug H, Maechler M, Bolker B. 2017 glmmTMB Balances Speed and flexibility among packages for zero-inflated generalized linear mixed modeling. R J 9, 378–400. (doi:doi:10.32614/RJ-2017-066.)

47. Lüdecke D, Ben-Shachar M, Patil I, Waggone P, Makowski D. 2021 “performance: An R Package for Assessment, Comparison and Testing of Statistical Models. J Open Source Softw 6, 3139. (doi:doi:10.21105/joss.03139.)

48. Chung Y, Rabe-Hesketh S, Dorie V, Gelman A, Liu J. 2013 A nondegenerate penalized likelihood estimator for variance parameters in multilevel models. Psychometrika 78, 685–709. (doi:10.1007/S11336-013-9328-2/FIGURES/9)

49. Fox J, Weisberg S. 2019 An R Companion to Applied Regression, Third Edition. Thousand Oaks CA: Sage.

50. In press. Lenth R (2024). emmeans: Estimated Marginal Means, aka Least-Squares Means_. R package version 1.10.0, <https://CRAN.R-project.org/package=emmeans>. (doi:10.1080/00031305.1980.10483031)

51. Lefcheck JS. 2016 piecewiseSEM: Piecewise structural equation modelling in r for ecology, evolution, and systematics. Methods Ecol Evol 7, 573–579. (doi:10.1111/2041-210X.12512)

52. Lüdecke D, Ben-Shachar MS, Patil I, Waggoner P, Makowski D. In press. performance: An R Package for Assessment, Comparison and Testing of Statistical Models. (doi:10.21105/joss.03139)

53. Chung DJ, Bryant HJ, Schulte PM. 2017 Thermal acclimation and subspecies-specific effects on heart and brain mitochondrial performance in a eurythermal teleost (Fundulus heteroclitus). Journal of Experimental Biology 220, 1459–1471. (doi:10.1242/JEB.151217/262557/AM/THERMAL-ACCLIMATION-AND-SUBSPECIES-SPECIFIC)

54. Scott KY, Matthew R, Woolcock J, Silva M, Lemieux H. 2019 Adjustments in the control of mitochondrial respiratory capacity to tolerate temperature fluctuations. Journal of Experimental Biology 222. (doi:10.1242/JEB.207951/267297/AM/ADJUSTMENTS-IN-CONTROL-OF-MITOCHONDRIAL)

55. De Bonville J et al. 2025 Dynamics of thermal tolerance plasticity across fish species and life stages. J Therm Biol 127, 104024. (doi:10.1016/J.JTHERBIO.2024.104024)

56. Chung DJ, Schulte PM. 2020 Mitochondria and the thermal limits of ectotherms. Journal of Experimental Biology 223. (doi:10.1242/JEB.227801/226043)

57. López-Otín C, Blasco MA, Partridge L, Serrano M, Kroemer G. 2023 Hallmarks of aging: An expanding universe. Cell 186, 243–278. (doi:10.1016/J.CELL.2022.11.001)

58. Ishikawa A, Kitano J, Dickinson MH, Vosshall LB, Dow JAT. 2020 Diversity in reproductive seasonality in the three-spined stickleback, Gasterosteus aculeatus. Journal of Experimental Biology 223. (doi:10.1242/JEB.208975)

