## Supplementary Material for "Seasonal decline in mitochondrial plasticity in the three-spined stickleback"

*for*

### A. SUPPLEMENTARY METHODS

*Study species and fish husbandry*

Pre-reproductive adult three-spined sticklebacks (*Gasterosteus aculeatus*) were collected in early March 2024 from the River Kelvin (55°52'29.1"N, 4°16'48.5"W), Glasgow, Scotland, UK, using dip nets and minnow traps. Adult sticklebacks typically breed in shallow water, where temperatures can fluctuate by as much as 10-18 °C within a single day [1,2]. Stream populations at mid-latitudes (50–60° N), such as those in Britain, generally reproduce for 3.5–4 months during spring and summer (April–August) [3].

Captured fish were transferred into 18-litre plastic buckets containing river water aerated using battery-powered air pumps. A total of 200 individuals were transported to the laboratory within 2 hours. Upon arrival, fish were gradually acclimatized to laboratory conditions by slowly replacing river water with aquarium water over approximately 1 hour. No mortality occurred during transport or acclimatization. To minimize handling stress, fish were not measured at this stage. Following this initial settling period, fish were housed in groups in 80-litre fiberglass holding tanks, each containing shelter structures (rocks and plastic plants) and supplied with recirculating UV-sterilized, aerated water maintained at 12°C (the approximate water temperature of Scottish rivers in the breeding season; see below). The photoperiod was fixed at 12:12 (light:dark) to exclude seasonal variation in daylight exposure which could confound the effects of age.

To prevent aggression and reduce social stress, any fish that developed male secondary sexual characteristics (i.e., blue coloration of eyes and/or red coloration of throat; [4]) were removed from the tanks so that the tanks only contained female or nonreproductive male fish. Even though the females were kept without breeding age males, they can lay clutches of eggs and cannot easily be prevented from so doing. Any eggs laid were unfertilized and were not reared; they either settled in the tank or degraded over time. They could not be attributed to individual females. Consequently, it was not possible to have an end-of-season group that had not produced any eggs, and variation in egg production could potentially have contributed to variation in the state of females late in the season. However, any energetic costs of egg production were likely to be minor, as females were provided with ad libitum food and did not rear offspring. Fish were fed ad libitum once daily with commercial pellet food.

*Tissue sampling and mitochondria isolation*

Mitochondria were isolated from white muscle tissue following euthanasia via a benzocaine overdose (10 mg/L). Isolates of muscle mitochondria were prepared using a modified version of previous protocols [5]. Immediately after death, muscle samples from the myotome were dissected and placed on ice in isolation buffer (100 mM sucrose, 50 mM Tris base, 5 mM $MgCl_2$, 5 mM EGTA, 100 mM KCl, 1 mM ATP; pH 7.1). Muscle extracts were minced and gently homogenized using a Teflon-on-glass homogenizer with six strokes at 100 rpm. The resulting homogenate was centrifuged at 1000 g (10 min). The supernatant was then centrifuged at 8700 g (10 min). Resulting pellets were resuspended in in 1000 µl of storage buffer (0.5 mM EGTA, 3 mM $MgCl_2$, 60 mM potassium methane sulphonate, 20 mM taurine, 10 mM $KH_2PO_4$, 20 mM HEPES, 110 mM sucrose, 0.02 mM vitamin E succinate, 2 mM pyruvate, and 2 mM malate; pH 7.1). A portion of the mitochondrial preparation was kept on ice for immediate mitochondrial function measurements, and the remainder was stored at −80 °C for subsequent enzymatic analysis.

*Mitochondrial function measurements*

Mitochondrial function was assessed at 21 °C using high-resolution respirometry on Oxygraph-2k systems (Oroboros Instruments, Innsbruck, Austria). For the fish kept continuously at 12 °C, mitochondrial respiration was measured at both 12 °C and 21 °C to additionally evaluate whether assay temperature influenced potential age-related effects on mitochondrial function. Isolated mitochondrial samples were introduced into the respirometry chamber containing 2 ml of MiR05 respiration buffer

($0.5$ mM EGTA, 3 mM $MgCl_2$, 60 mM potassium lactobionate, 20 mM taurine, 10 mM $KH_2PO_4$, 20 mM HEPES, 110 mM sucrose, and 1 g/l fatty acid-free BSA, adjusted to pH 7.3). Oxygen consumption was determined by monitoring the rate of oxygen decline in the sealed chamber. Mitochondrial respiration was initially measured in the absence of ADP following the addition of malate (2 mM) and pyruvate (5 mM). Subsequently, a saturating concentration of ADP (5 mM) was added to activate full oxidative phosphorylation (OXPHOS) through complex I in the presence of pyruvate and malate. Next, glutamate (10 mM) followed by succinate (25 mM) were added to evaluate maximal OXPHOS capacity via complexes I and II (maximum OXPHOS). This rate of respiration in the presence of all substrates represents the maximum rate of OXPHOS, since it is occurring under conditions where both Complex I and Complex II can operate with an excess of substrates and ADP. Thus, OXPHOS can be used as a proxy of the capacity for mitochondrial ATP production. To control for the permeability of the outer mitochondrial membrane, exogenous cytochrome c (10 μM) was added, which caused only a relatively low increase in respiration (18.3% on average), indicating that mitochondrial preparations were of high quality. A small subset of samples was excluded due to technical issues or compromised mitochondrial quality (i.e., outer membrane permeability; Table S1 summarizes the number of samples at each experimental phase). Oligomycin was then used to determine LEAK respiration (the oxygen consumption that is used to offset the leakage of protons across the inner mitochondrial membrane rather than to generate ATP). Non-mitochondrial oxygen consumption was quantified by adding antimycin A and subsequently subtracted from all respiration measurements. Mitochondrial respiration was normalized to citrate synthase activity, a widely used biomarker for mitochondrial volume [6,7], allowing us to control for mitochondrial content (see below).

**Table S1.** Number of individuals at each phase of the study. "Sampled" refers to individuals used for mitochondrial function measurements; "Analysed" refers to individuals retained after quality control (see Methods for details).

| Age | Temperature treatment | Tank (n) | Sampled | Analysed |
|---|---|---|---|---|
| Young | Constant cool 12 °C | 21 | 18 | 9 |
| | Diel cycling 12-21 °C | 21 | 20 | 9 |
| | Constant warm 21 °C | 21 | 19 | 10 |
| Old | Constant cool 12 °C | 21 | 18 | 14 |
| | Diel cycling 12-21 °C | 21 | 20 | 17 |
| | Constant warm 21 °C | 21 | 20 | 18 |

OXPHOS Coupling Efficiency (OxCE) was calculated as previously described in [8]:

$$OxCE = 1 - (LEAK / OXPHOS)$$

This mitochondrial efficiency index ranges from 0 (indicating no ATP production despite substrate availability) to 1 (indicating absence of leak respiration and all oxygen consumption dedicated to ATP synthesis). Thus, respiration rates indicate potential metabolic capacity levels, while OxCE indicates how efficient mitochondria are at using oxygen to produce ATP [8].

*Citrate synthase activity*

At the end of the respirometry assay, we recovered isolated mitochondrial subsamples from the O2k chamber and stored these -80 C after for subsequent determination of citrate synthase (CS) activity. Citrate synthase (CS) activity was measured for all samples under identical, temperature-controlled conditions using a SpectraMax Plus 384 spectrophotometer (Molecular Devices), following an adapted protocol [9]. While CS activity is temperature-sensitive, applying identical standardized assay across all samples ensures consistency and comparability. Isolated mitochondria were homogenized using a PowerGen 125 Homogenizer (Fisher Scientific) in a 1:1 dilution with homogenization buffer composed of 100 mM $KH_2PO_4$, 1 mM EGTA, 1 mM EDTA, 0.1% Triton X-100 (pH 7.2), and 1 mM phenylmethylsulphonyl fluoride (PMSF). The diluted homogenates were centrifuged at 1000 × g for 5 minutes at 4°C, and the supernatant was collected. CS activity was quantified in these supernatants by measuring the change in absorbance at 412 nm in triplicate over time. The assay mixture contained 100 mM $KH_2PO_4$ (pH 8.0), 0.15 mM acetyl-CoA, 0.15 mM 5,5'-dithiobis-(2-nitrobenzoic acid), and 0.5 mM oxaloacetate. Using an extinction coefficient (ε) of 14.15 $mM^{-1}$ $cm^{-1}$, enzyme activity was expressed as μmol of tissue per minute.

### B. SUPPLEMENTARY TABLES

**Table S2.** Summary of linear mixed models (LMMs) testing the effects of temperature treatment, age, and cytochrome c (Cyt C) on mitochondrial ATP-producing respiration (OXPHOS), non-ATP-producing respiration (LEAK), and oxidative coupling efficiency (OxCE). The marginal $R^2$ ($R^2_m$) indicates the proportion of variance explained by the fixed effects alone, while the conditional $R^2$ ($R^2c$) shows the variance explained by both fixed and random effects. Statistically significant terms are highlighted in bold.

| OXPHOS | Chi sq | df | p-value |
|---|---|---|---|
| Temperature treatment | 1.481 | 2 | 0.477 |
| Age | 4.465 | 1 | **0.035** |
| Cyt C | 0.074 | 1 | 0.786 |
| Temperature treatment x Age | 23.916 | 2 | **< 0.001** |

| Random | Variance | Std. Dev. |
|---|---|---|
| Tank ID | 0.247 | 0.497 |
| Chamber ID | 0.069 | 0.263 |
| Residual | 1.199 | 1.095 |
| Conditional $R^2$ | 0.435 | |
| Marginal $R^2$ | 0.287 | |

| LEAK | Chi sq | df | p-value |
|---|---|---|---|
| Temperature treatment | 1.557 | 2 | 0.459 |
| Age | 0.193 | 1 | 0.661 |
| Cyt C | 11.572 | 1 | **< 0.001** |
| Temperature treatment x Age | 23.825 | 2 | **< 0.0001** |

| Random | Variance | Std. Dev. |
|---|---|---|
| Tank ID | 0.009 | 0.093 |
| Chamber ID | 0.002 | 0.043 |
| Residual | 0.024 | 0.157 |
| Conditional $R^2$ | 0.516 | |
| Marginal $R^2$ | 0.310 | |

| OxCE | Chi sq | df | p-value |
|---|---|---|---|
| Temperature treatment | 0.011 | 2 | 0.994 |
| Age | 0.372 | 1 | 0.542 |
| Cyt C | 0.354 | 1 | 0.552 |

| Random | Variance | Std. Dev. |
|---|---|---|
| Tank ID | 0.074 | 0.272 |
| Chamber ID | 0.080 | 0.283 |
| Conditional $R^2$ | 0.067 | |
| Marginal $R^2$ | 0.024 | |

**Table S3.** Results of the structural equation model (SEM) testing the effects of temperature treatment and cytochrome c (a quality-control for outer membrane permeability; CytC) on mitochondrial ATP-producing respiration (OXPHOS), and the downstream effects on body condition. Significant effects ($p < 0.05$) are in bold. Model fit was tested using the Fisher's C, with $p > 0.05$ indicating no significant missing paths.

| Response | Predictor | Estimate | Std. Error | df | Crit. Value | *p*-value |
|---|---|---|---|---|---|---|
| OXPHOS | Temperature treatment | - | - | 2 | 10.817 | **0.005** |
| OXPHOS | Warm 21C | -0.476 | 0.321 | 21 | -1.482 | 0.153 |
| OXPHOS | Diel cycling 12-21C | -0.181 | 0.339 | 21 | -0.534 | 0.599 |
| OXPHOS | Cool 12C | 0.876 | 0.317 | 21 | 2.763 | **0.012** |
| OXPHOS | CytC | 0.231 | 0.164 | 28 | 1.412 | 0.158 |
| Body Condition | OXPHOS | -0.064 | 0.186 | 28 | -0.346 | 0.729 |
| Body Condition | Temperature treatment | - | - | 2 | 13.814 | **0.001** |
| Body Condition | Warm 21C | -0.831 | 0.267 | 22 | -3.111 | **0.005** |
| Body Condition | Diel cycling 12-21C | 0.284 | 0.259 | 22 | 1.094 | 0.286 |
| Body Condition | Cool 12C | 0.639 | 0.299 | 22 | 2.138 | **0.044** |
| Global goodness-of-fit: Fisher's C = 2.75; *p*-value = 0.253 | | | | | | |

**Table S4.** Results of the structural equation model (SEM) testing the effects of temperature treatment and cytochrome c (a quality-control for outer membrane permeability; CytC) on mitochondrial non-ATP-producing respiration (LEAK), and the downstream effects on body condition. Significant effects ($p < 0.05$) are in bold. Model fit was tested using the Fisher's C, with $p > 0.05$ indicating no significant missing paths.

| Response | Predictor | Estimate | Std. Error | df | Crit. Value | *p*-value |
|---|---|---|---|---|---|---|
| LEAK | Thermal regime | - | - | 2 | 6.280 | **0.043** |
| LEAK | Warm 21C | -0.29 | 0.331 | 21 | -0.883 | 0.387 |
| LEAK | Diel cycling 12-21C | -0.215 | 0.354 | 21 | -0.607 | 0.550 |
| LEAK | Cool 12C | 0.732 | 0.333 | 21 | 2.199 | **0.039** |
| LEAK | CytC | 0.481 | 0.154 | 28 | 3.125 | **0.002** |
| Body Condition | LEAK | -0.057 | 0.167 | 28 | -0.344 | 0.731 |
| Body Condition | Thermal regime | - | - | 2 | 15.464 | **< 0.001** |
| Body Condition | Warm 21C | -0.820 | 0.256 | 22 | -3.207 | **0.004** |
| Body Condition | Diel cycling 12-21C | 0.288 | 0.259 | 22 | 1.113 | 0.278 |
| Body Condition | Cool 12C | 0.623 | 0.278 | 22 | 2.241 | **0.036** |
| Global goodness-of-fit: Fisher's C = 2.91; *p*-value = 0.233 | | | | | | |

### C. SUPPLEMENTARY FIGURES

Each tank in the thermal acclimation trials was equipped with air stones and shelter structures (rocks and plastic plants), and water flow was maintained continuously on a recirculation system: approximately 24 L/hour for heated tanks and 12–16 L/hour for tanks held at 12°C.

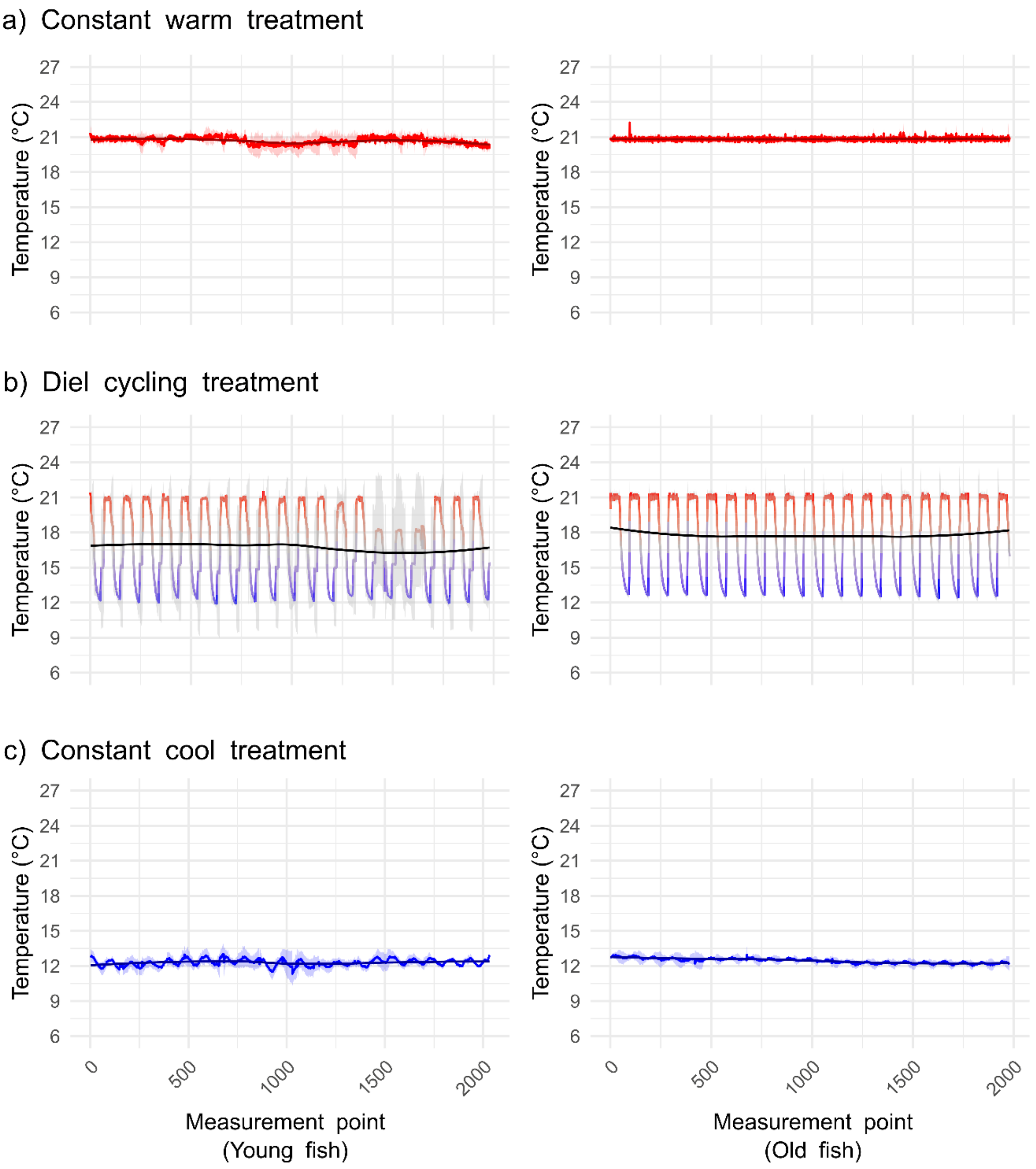


**Figure S1. Temperature conditions in the experimental tanks during the 21-day exposure period for the three temperature treatments: (a) constant cool (12 °C), (b) diel cycling (12–21 °C), and (c) constant warm (21 °C).** Water temperature in the thermal acclimation tanks was recorded using HOBO loggers every 15 min using temperature sensors (measurement points). For each treatment, the dark line represents the mean temperature across three replicate tanks (± standard deviation; shaded area). The black solid line shows a locally smoothed trend in the mean temperature for visualization. Across all tanks, mean (± SD) temperatures were: constant cool = 12.3 ± 0.5 °C (young fish) and 12.4 ± 0.3 °C (old fish); diel cycling = 16.7 ± 3.9 °C (young fish) and 17.6 ± 3.6 °C (old fish); and constant warm = 20.6 ± 0.5 °C (young fish) and 20.8 ± 0.2 °C (old fish).

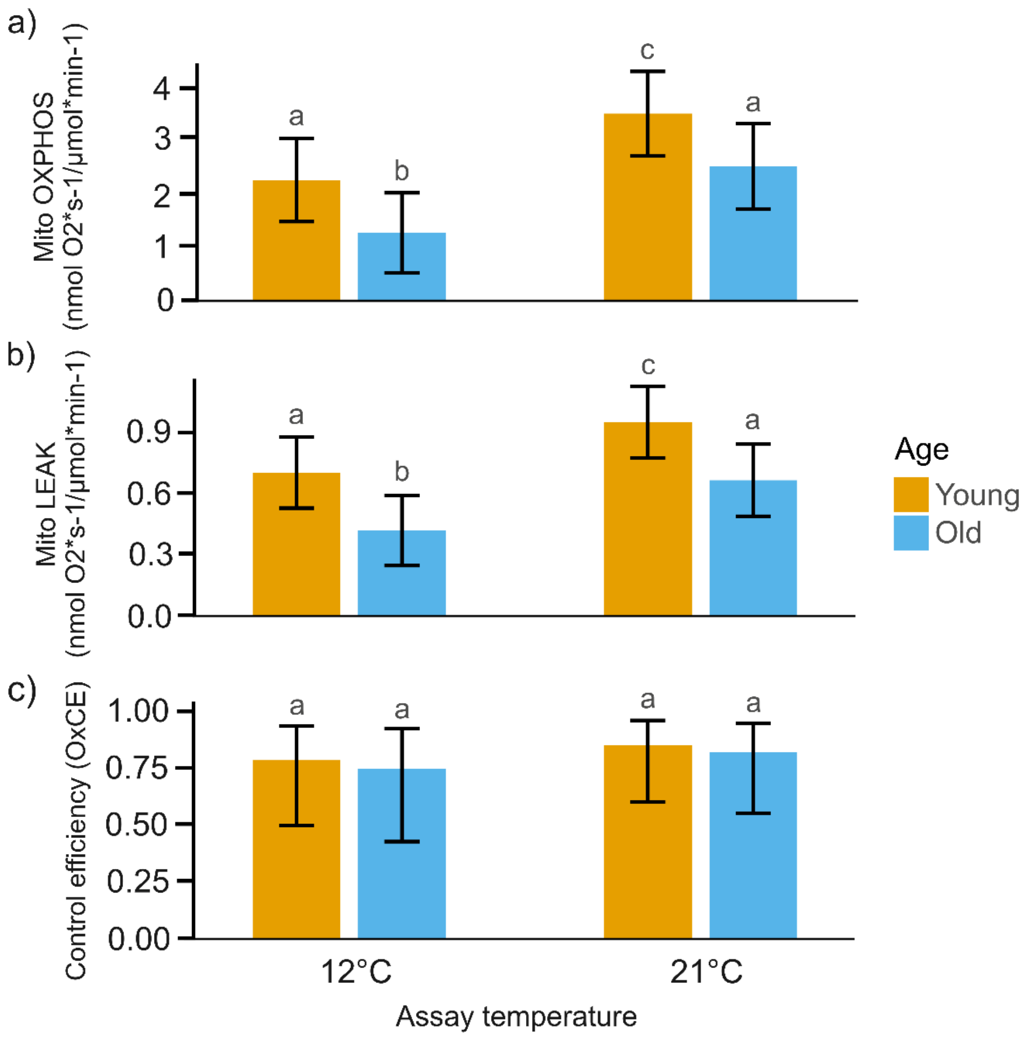


**Figure S2: Mitochondrial plasticity in response to assay temperature is age-dependent in cool-acclimated fish.** Isolated mitochondria of "young" and "old" three-spined sticklebacks acclimated to 12°C were assayed for (a) OXPHOS respiration, (b) LEAK respiration and (c) oxidative control efficiency (OxCE) at both 12°C and 21°C. Values are shown as estimated marginal means ± 95% CI. Different letters indicate significant differences between groups (Tukey's HSD, $p < 0.05$).

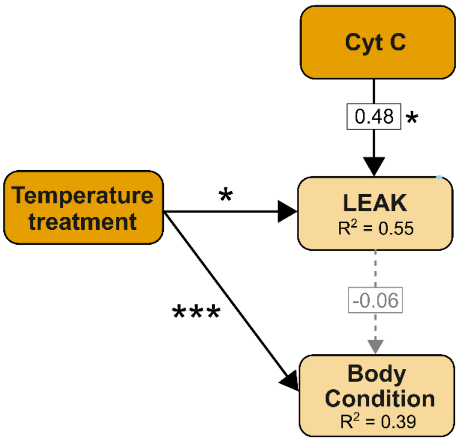


**Figure S3. Direct and indirect paths between thermal regime and body condition (BCI).** Indirect paths included non-ATP-producing respiration (LEAK). Grey dotted arrows represent non-significant paths, while black solid lines represent significant paths (** $p$ < 0.01). $R^2$ represents the proportion of variance explained in response variables by a set of predictors. Standardized estimates are reported for those paths including continuous variables. The model output is shown in Table S3.

### REFERENCES


1. Corey E, Linnansaari T, Cunjak RA, Currie S. 2017 Physiological effects of environmentally relevant, multi-day thermal stress on wild juvenile Atlantic salmon (Salmo salar). *Conserv Physiol* **5**, cox014. (doi:10.1093/CONPHYS/COX014)

2. Broadmeadow SB, Jones JG, Langford TEL, Shaw PJ, Nisbet TR. 2011 The influence of riparian shade on lowland stream water temperatures in southern England and their viability for brown trout. *River Res Appl* **27**, 226–237. (doi:10.1002/RRA.1354)

3. Ishikawa A, Kitano J. 2020 Diversity in reproductive seasonality in the three-spined stickleback, Gasterosteus aculeatus. *Journal of Experimental Biology* **223**. (doi:10.1242/JEB.208975,)

4. Wooton R. 1978 *The Biology of the Sticklebacks*. Academic Press. (doi:10.1002/IROH.19780630314)

5. Dawson NJ, Scott GR. 2022 Adaptive increases in respiratory capacity and O2 affinity of subsarcolemmal mitochondria from skeletal muscle of high-altitude deer mice. *The FASEB Journal* **36**, e22391. (doi:10.1096/FJ.202200219R)

6. McLaughlin KL, Hagen JT, Coalson HS, Nelson MAM, Kew KA, Wooten AR, Fisher-Wellman KH. 2020 Novel approach to quantify mitochondrial content and intrinsic bioenergetic efficiency across organs. *Sci Rep* **10**, 1–15. (doi:10.1038/S41598-020-74718-1;SUBJMETA=1465,319,333,443,45,475,631;KWRD=ENERGY+METABOLISM,PROTEOMICS)

7. Larsen S *et al.* 2012 Biomarkers of mitochondrial content in skeletal muscle of healthy young human subjects. *J Physiol* **590**, 3349–3360. (doi:10.1113/JPHYSIOL.2012.230185)

8. Koch RE *et al.* 2021 Integrating mitochondrial aerobic metabolism into ecology and evolution. *Trends Ecol Evol* **36**, 321–332. (doi:10.1016/J.TREE.2020.12.006/ASSET/0EBF99B9-96F2-40DC-B316-EF6FE592DB20/MAIN.ASSETS/MMC2.MP4)

9. Dawson NJ, Lyons SA, Henry DA, Scott GR. 2018 Effects of chronic hypoxia on diaphragm function in deer mice native to high altitude. *Acta Physiologica* **223**, e13030. (doi:10.1111/APHA.13030;WGROUP:STRING:PUBLICATION)
